# iterativeWGCNA: iterative refinement to improve module detection from WGCNA co-expression networks

**DOI:** 10.1101/234062

**Authors:** Emily Greenfest-Allen, Jean-Philippe Cartailler, Mark A. Magnuson, Christian J. Stoeckert

## Abstract

Weighted-gene correlation network analysis (WGCNA) is frequently used to identify highly co-expressed clusters of genes (modules) within whole-transcriptome datasets. However, transcriptome-scale networks tend to be highly connected, making it challenging for the hierarchical clustering underlying the WGCNA-based classification to discriminate coherently expressed gene sets without significant information loss from either *a priori* filtering of the expression dataset or *a posteriori* pruning of the cluster dendrogram.

Here we present iterativeWGCNA, a Python-wrapped extension for the WGCNA R software package that improves the robustness of detected modules and minimizes information loss. The method works by pruning poorly fitting genes from estimated modules and then re-running WGCNA to refine gene clusters. After refining, pruned genes are assembled into a new expression dataset to isolate overlapping modules and the process repeated. In doing so, iterativeWGCNA provides an unsupervised, non-biased filtering to generate a robust, comprehensive network-based classification of whole-transcriptome expression datasets.

## Background

Weighted Gene Correlation Network Analysis (WGCNA) is a widely used method for classifying genes via hierarchical clustering of gene co-expression networks (GCNs). GCNs are a particularly useful for analyzing high-dimensionality expression datasets as they provide an intuitive framework for describing changes in expression driven by cellular processes acting within or across multiple conditions (Langfelder and Horvath, 2008). Consequently, WGCNA is now commonly used to analyze whole transcriptome expression datasets, obtained from both microarray and RNA-seq expression analysis.

WGCNA is very much an all-in-one toolbox for GCN analysis. Using WGCNA, a GCN is constructed from a weighted adjacency matrix defining the connection strength among gene pairs. Connection strength is determined as pairwise-correlation between gene expression profiles across samples, adjusted by a power-law function that down-weights weaker correlations such that the GCN approaches a scale-free topology. This in turn is used to estimate the topological overlap among genes (adjacency adjusted for the proportion of shared connections), and the topological overlap-based dissimilarity matrix is subject to hierarchical clustering. Finally, by cutting the cluster dendrogram at a height that maximizes the intra-connectedness among cluster genes, WGCNA organizes genes into co-expressed gene sets. Ideally these clusters should approximate true network modules, in that member genes are expected to be highly intra-connected, but sparsely connected to all other such groups.

The effectiveness of this approach depends on the assumption that the GCN is scale-free and that it has an inherent, hierarchical structure (i.e., little overlap between modules). A scalefree network is self-organized such that the number of links or degree, *k*, originating from a node is directly proportional to the probability of a connection to a node, *P*(*k*), following a power-law distribution. The scale-free nature of a network can be approximated as the slope of a log-log plot of the network connectivity distribution, which is expect to approach 1.0 (Zhang and Horvath, 2005). The soft-thresholding power in WGCNA is selected to maximize that fit for the expression dataset and is key to effective WGCNA analysis. However, for whole-transcriptome datasets, the scale-free topology fit index (signed *r*^2^ to the fit) may fail to reach values >0.8 for reasonable powers (<30 for a signed network) (Horvath, 2017). Consequently, WGCNA-based classifications of transcriptome-scale expression datasets tend to result in gene clusters too poorly resolved to establish a robust framework without extensive pruning.

Transcriptome-scale analyses tend to be either inherently heterogenous (conditions are broadly defined) or extremely homogenous (e.g., single-cell sequencing, cell maturation). In both cases, a subset of the samples may be globally different from the rest (e.g., distinct cells or tissues in the former; genes defining cell identity versus those specifying cellular process, the latter) or strongly correlated only across subsets of conditions (e.g., time series), all of which can yield significant correlations among large groups of genes. In these cases, most genes have high degree; the GCNs are highly connected and consequently lack both hierarchical organization and scale-free topology that cannot be approximated even after down-weighting weaker correlations via power-law scaling.

High connectivity creates a large amount of overlap in gene-neighborhoods, making it challenging for greedy hierarchical clustering to partition genes into discrete, non-overlapping clusters. This is readily seen in the poor resolution of the hierarchical clustering figured in many published whole-transcriptome WGCNA studies (e.g., (Xue et al., 2013), which are typically displayed as a dendrogram illustrating the hierarchical clustering of the gene set, paired with a color scale indicating the module membership of each gene (**Fig. 1D,F**). Simple visual inspection of these dendrograms is sufficient to highlight a poorly resolved result, which is characterized by weakly segregated clusters (separated by short branch lengths) that group genes across a spectrum of weakly to strongly correlated expression profiles (long branch lengths within clusters and colors identifying module membership are distributed across disparate gene-clusters). Consequently, the use of WGCNA with whole-transcriptome datasets has generally been *a priori* limited to a selected subset of expressed genes (Massa et al., 2011), or has focused on the central core of the best defined modules (Xue et al., 2013). In either case, genes lacking coherent expression across the samples are downplayed or disregarded.

**Figure 1.**
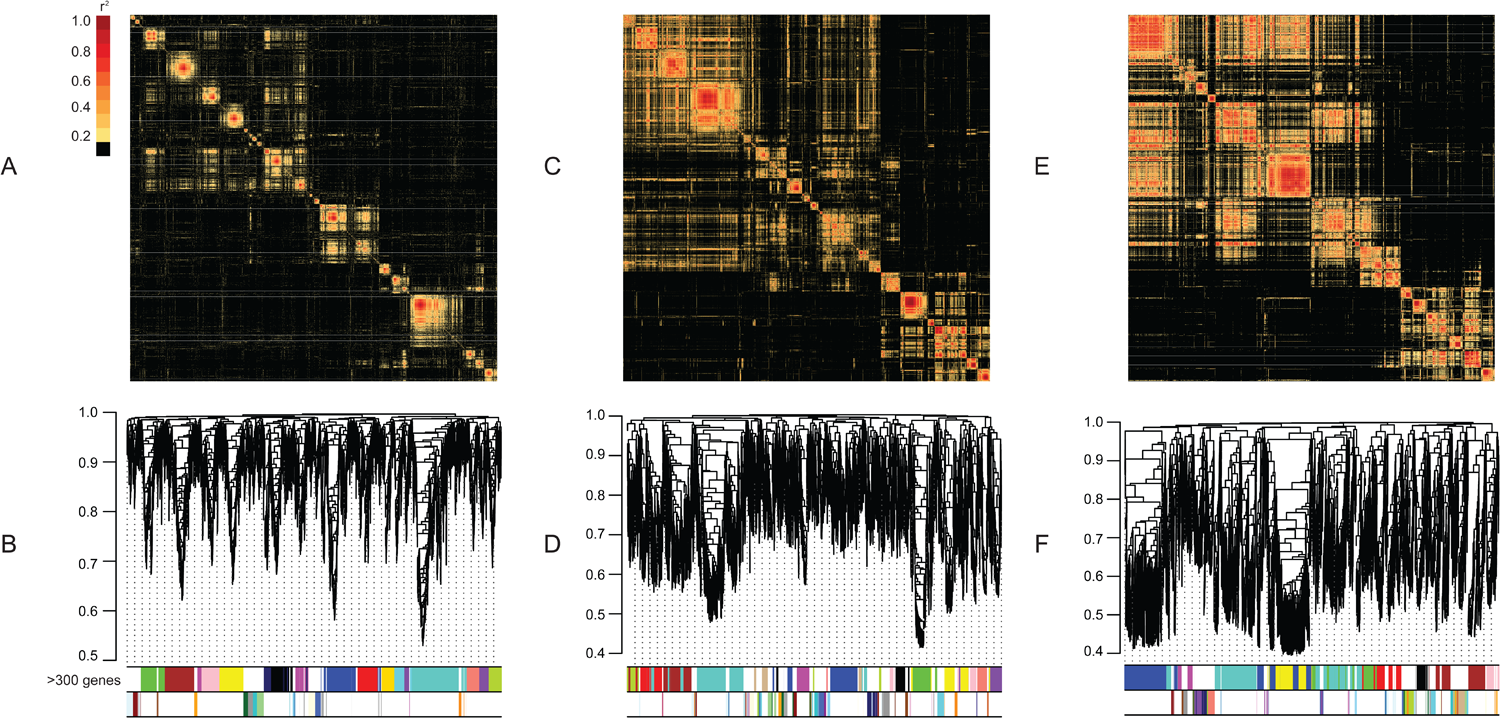
Network block models and WGCNA-based clustering of GCN from idealized and real datasets. A. Block model of the weighted-adjacency matrix defining the GCN of the idealized simulated dataset (SIM; see methods) illustrates its hierarchically organized topology. Strongest connections between gene-pairs (red, dark orange) are concentrated along the matrix diagonal delimitating a series of well-defined “blocks” corresponding to robust network modules. Few inter-modular connections exist. Ordering of genes same as in B, below. B. Clustering and module assignment from standard WGCNA analysis of the GCN illustrated in A, above. The colored strips under each dendrogram depict the module assignment for the gene associated with the corresponding dendrogram branch. To highlight the module cores and the correspondence between the network block model (A, above) and dendrogram clustering, modules are split into groups two based upon size: upper strip: modules containing ≥300 genes; lower strip: modules containing <300 genes. Module colors are not conserved among panels. C. Block model of the weighted-adjacency matrix defining the GCN of the ADULT dataset, illustrating a visible central core along the diagonal that is muddled by both network substructure (small blocks inside of big blocks) and numerous inter-modular connections. Coloring as in A, above. Ordering of genes same as in D, below. D. Clustering and module assignment from standard WGCNA analysis of the GCN illustrated in C, above. The higher inter-modular connectivity of this network, compared to the simulated dataset (A) is sufficient to negatively affect the robustness of the hierarchical clustering. Robust modules corresponding to the best-defined blocks (C above) can be recognized, but overall branch lengths are longer and the detected modules not as well-delimited by the dendrogram structure. E. Block model of the weighted-adjacency matrix defining the GCN of the FETAL dataset, illustrating a lack of hierarchical organization except at the broadest scales, reflecting the noisier nature of the dataset. Module blocks are larger, and not as well defined; with some intermodular connections being very strong (red). Coloring as in A, above. Ordering of genes same as in D, below. F. Clustering and module assignment from standard WGCNA analysis of the GCN illustrated in E, above. With few exceptions (e.g., black and pink modules), the hierarchical clustering is unable to robustly discriminate co-expressed gene sets. Long branch lengths within geneclusters indicate groupings of genes with low topological overlap (and therefore very dissimilar expression profiles). Detected modules are not well-delimited by the dendrogram structure.

Here we introduce an iterative extension to WGCNA to improve the robustness of whole-transcriptome GCN clustering and reduce the information loss created by filtering or disregarding poorly resolved clusters. This method builds on WGCNA by pruning poorly fitting genes from estimated modules and then re-running WGCNA to generate refined, coherent clusters of highly co-expressed genes. After refining, residual genes (those pruned through all iterations) are assembled into a new expression dataset to isolate overlapping modules and the process repeated. In doing so, our approach utilizes an unsupervised, non-biased filtering to generate a robust network-based classification of whole-transcriptome datasets.

## Method

### Iterative application of Weighted Gene Correlation Network Analysis (iterativeWGCNA)

The iterativeWGCNA approach is outlined in **Fig. 2**. The process consists of five steps: 1) module detection, 2) goodness of fit evaluation, 3) network refinement, 4) residual network estimation, and 5) network assembly. A single iteration of the method comprises steps 1-3, which use the *blockwiseModules* function in the WGCNA R package (Langfelder and Horvath, 2008) to efficiently infer and cluster a co-expression network from the expression dataset (step 1), calculate eigengenes (representative expression profile for each module; its first principal component) and evaluate the goodness of fit using eigengene connectivity (*k_ME_*) of each gene to its assigned module (step 2). Eigengene connectivity is the correlation between a gene’s expression profile and the module eigengene (Langfelder and Horvath, 2008). The network is refined (step 3) by recalculating the goodness of fit using the WGCNA *corAndPvalue* function and removing any retained genes classified by the *blockwiseModules* function that have poor eigengene connectivity to their assigned module (*k_ME_* < minimum *k_ME_* to stay; *p*-value < reassignment threshold). A new iteration is then initiated by running the *blockwiseModules* function on the reduced expression dataset comprised only of retained, well-classified genes. Steps 1-3 are repeated until no further genes are pruned, completing a single pass of the refinement algorithm over the dataset.

**Figure 2.**
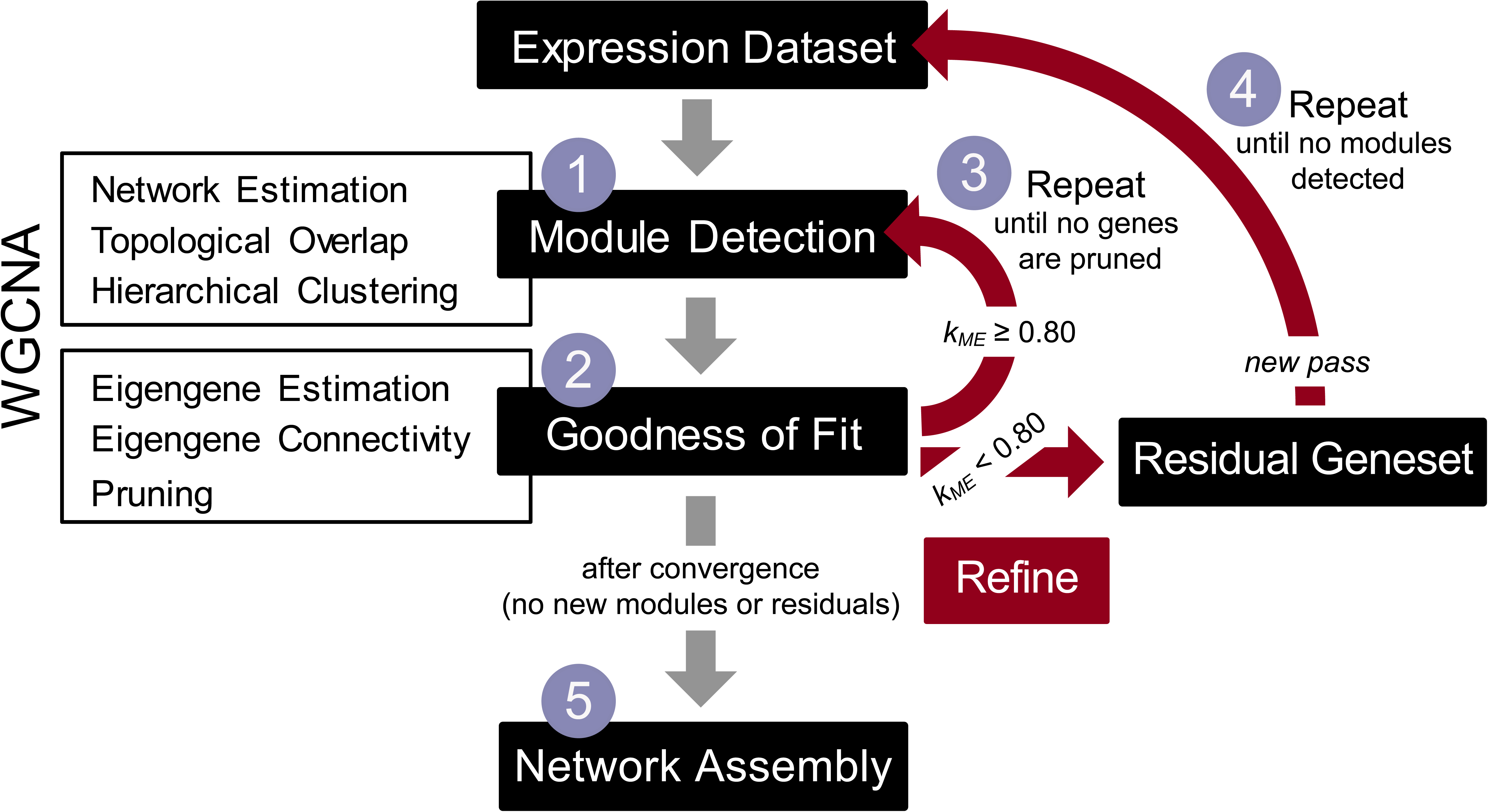
Flow chart illustrating iterative refinements (red) to blockwise WGCNA allowing the inference of increasingly modular networks and isolation of highly co-expressed gene clusters. Numbered blue circles highlight steps referenced in **Fig. 4** and below. A single iteration of this method comprises steps 1 and 2. In step 1, WGCNA’s *blockwiseModules* function is used to efficiently infer and cluster a co-expression network from the complete expression dataset. In step 2, blockwise WGCNA is used to calculated module eigengenes and evaluate the goodness of fit of each gene to its assigned module using eigengene connectivity (*k_ME_*). Poorly fitting genes are pruned, and another iteration (step 3) is run by applying blockwise WGCNA to the smaller expression dataset comprised only of retained, well-classified genes. Steps 1-3 are repeated until no further genes were pruned, completing a single pass of the refinement algorithm over the dataset. Subsequent passes (step 4) are initiated by assembling residual genes (those pruned through all iterations) into a new expression dataset. Steps 1-4 are repeated until no additional modules are isolated. After full convergence (no new modules or residuals to the fit), the network is assembled (step 5) by evaluating the goodness of fit of all genes to all predicted modules. Genes are assigned to the module to which they showed the best fit; genes exhibiting poor fit to all module eigengenes are left unclassified.

Subsequent passes (step 4) are initiated by assembling residual genes (those pruned through all iterations of the previous passes) into a new expression dataset. Steps 1-4 are repeated until a second convergence condition, that no additional modules are detected, is met. After convergence, the network is assembled (step 5) by evaluating the goodness of fit of all genes to all predicted modules to account for the greedy nature of hierarchical clustering by allowing gene memberships to shift if a better fit was found to a module detected in a later pass. Genes are assigned to the module to which they have the best fit; genes with poor eigengene connectivity to all detected modules are left unclassified.

IterativeWGCNA was implemented for Python 3+, using the RPy2 Python-to-R interface to run embedded R code (Gautier, 2012, p. 2). The standalone Python module can be run on any platform supporting an installation of the R language and environment for statistical computing (version 3.0 or higher; R Core Development Team, 2017) and the WGCNA package for R (Langfelder and Horvath, 2008). It is available for download from GitHub (https://github.com/cstoeckert/iterativeWGCNA) and is registered in the Python Package Index (PyPI), enabling it to be installed via the Python package manager *pip*. Parameters for iterativeWGCNA are the same as for the *blockwiseModules* function in the WGCNA R package.

### Data Sources

We assessed the ability of iterativeWGCNA to improve the derivation of GCNs by using it to analyze two published datasets, as described below:

#### 1. Murine adult definitive red blood cells (ADULT)

This dataset is one of three microarray expression datasets generated to compare global gene expression during primitive, fetal definitive, and adult definitive erythropoiesis (Kingsley et al., 2013). The dataset comprises five high quality replicates of 4 maturational stages tracking murine adult definitive erythropoiesis from the proerythroblast thru reticulocyte stage isolated from bone marrow. The raw data were obtained from ArrayExpress (E-MTAB-1035). Normalization of probe expression data was performed using the *justGCRMA* function of the *gcrma* R package (Wu and Gentry, 2017). Gene expression was determined as the average probe expression across probes called present by MAS5, using the *mas5* function of the *affy* R package (Gautier et al., 2004), in ≥3 replicates of at least one of the 4 maturational stages. 12,937 genes were found to be expressed in this dataset.

#### 2. Murine fetal definitive red blood cells (FETAL)

Like the ADULT dataset, these data were assembled as part of a larger study comparing global gene expression during primitive, fetal definitive, and adult definitive erythropoiesis (Kingsley et al., 2013). This dataset comprises five replicates each of 4 maturational stages tracking murine fetal definitive erythropoiesis from the proerythroblast thru reticulocyte stage that are morphologically equivalent to those included in the ADULT dataset. Precursor erythroid cells were isolated from E14.5 liver; reticulocytes from E15.5 blood. Quality control analysis revealed that replicates for the nucleated erythroid precursor stages were not well differentiated, reflecting both rapid maturation during the expansion of the fetal erythroid niche and contamination from fetal liver cells (Kingsley et al., 2013). The raw data were obtained from ArrayExpress (E-MTAB-1035). Normalization and filtering were done as for the ADULT dataset. 12,794, genes were found to be expressed in this dataset.

### Simulated Data

An idealized, hierarchically organized, scale-free dataset (SIM) with similar magnitude and degree distribution to the murine blood cell datasets (20 conditions; 12,800 genes) was generated using the WGCNA function *simulateDatExpr*. The primary module structure was defined using the ADULT dataset as a template. WGCNA was applied to the ADULT dataset, resulting in 28 detected modules. The module eigengenes and proportion of genes assigned to each were passed to the *simulateDatExpr* function; all other parameters were set to minimize module overlap, noise, and secondary structure.

### Module Detection via iterativeWGCNA

WGCNA and its *blockwiseModules* function have many parameters that can be adjusted to affect the inference and clustering of a GCN. However, in practice most WGCNA-analyses are run using the default parameters, with only the power-law soft-threshold being specified depending on either the network type (6 for unsigned, 12 for signed), the complexity of the network (following the FAQ guidelines), or estimation using the *pickSoftThreshold* function in WGCNA. Here, we followed the industry standard for using the default parameter settings, performing iterativeWGCNA on each dataset, using the *blockwiseModules* default parameters for a “signed” network (power = 12).

### WGCNA Network Estimation and Module Detection

Standard WGCNA analysis was performed on each dataset following the steps outlined in the WGCNA tutorials and using recommended parameter settings for a signed network (power = 12; all others default) in the online FAQ as has become the standard in most studies applying WGCNA. WGCNA was used to calculate signed, power-law weighted adjacency matrices from the expression dataset and from those estimate a signed topological-overlap matrix (TOM). 1-TOM was used as a measure of dissimilarity and the matrix clustered using R’s *hclust* function. WGCNA was then used to determine module memberships by cutting the resulting dendrogram to maximize intra-modular connectivity, merge close modules (those with similar eigengenes), and plot the results.

Block models illustrating the topology of each GCN were generated by ordering the weighted adjacency matrices by the WGCNA clustering and then using the *pheatmap* R package to draw a heatmap.

## Results and Discussion

### Improved GCN partitioning with iterativeWGCNA

In whole-transcriptome expression datasets, genes can have many connections, and this is certainly the case for the FETAL dataset, where many genes have >500 connections even after power-law scaling. Despite this, subsets of these genes can define a hierarchically organized central “core” for the GCN, as illustrated by the red and orange squares of highly co-expressed genes flanking the diagonal of the network block models (**Fig. 1E**). The basic GCN clustering produced by WGCNA is driven by the strong signal established by these core gene sets and if they are well-isolated, WGCNA performs well, as in the case of the idealized simulated dataset (SIM), irrespective of the dimensionality (number of samples x number of genes) of the dataset (**Fig. 1A,B**). However, if that core-signal is blurred, the greedy nature of hierarchical clustering, which prioritizes locally optimal solutions, causes WGCNA to assign genes to the first best fit found among existing clusters, potentially assigning membership based on a weaker correlation (*r*^2^ ~ 0.8; orange, **Fig. 1**) instead of the global best fit (*r*^2^> 0.9; red; **Fig. 1**). Consequently, the best-defined modules are comprised of a central core of coherently expressed genes and a halo of genes that are not such a good fit.

As an alternative, using WGCNA’s *blockwiseModules* function improves the basic result by identifying and pruning genes that are a poor fit to their assigned module. With this algorithm, the goodness of fit of each gene to its module is assessed using the signed eigengene connectivity measure (*k_ME_*), or the correlation between a gene’s expression profile and the representative expression profile (eigengene) of the module (Langfelder and Horvath, 2008). Genes with an eigengene connectivity less than some selected *k_ME_* cutoff are considered a poor fit to their assigned module and “pruned” by WGCNA by assigning them to a “grey” module. Although this grey module is frequently a true “garbage bag”, including genes that can’t be unequivocally assigned to any module (e.g., those with expression profiles weakly correlated with multiple “core” profiles, are ubiquitously expressed, or exhibit oscillating or highly variable patterns of expression), it can also include genes assigned to the wrong module when prioritizing locally optimal solutions.

Pruning of poorly fitting genes typically improves the WGCNA result (**Fig. 3A**) compared to the non-pruned clustering (**Fig. 1C**), but the detected clusters can still be poorly resolved, especially in noisier datasets, such as the liver-contaminated FETAL dataset. In addition, pruning poorly classified alters the network architecture, in terms of both modularity and topological overlap, by reducing the number of inter-modular connections in the GCN. Thus, the measures of topological overlap used to drive the clustering, calculation of representative module eigengenes and justify the pruning, are no longer representative of the GCN.

**Figure 3.**
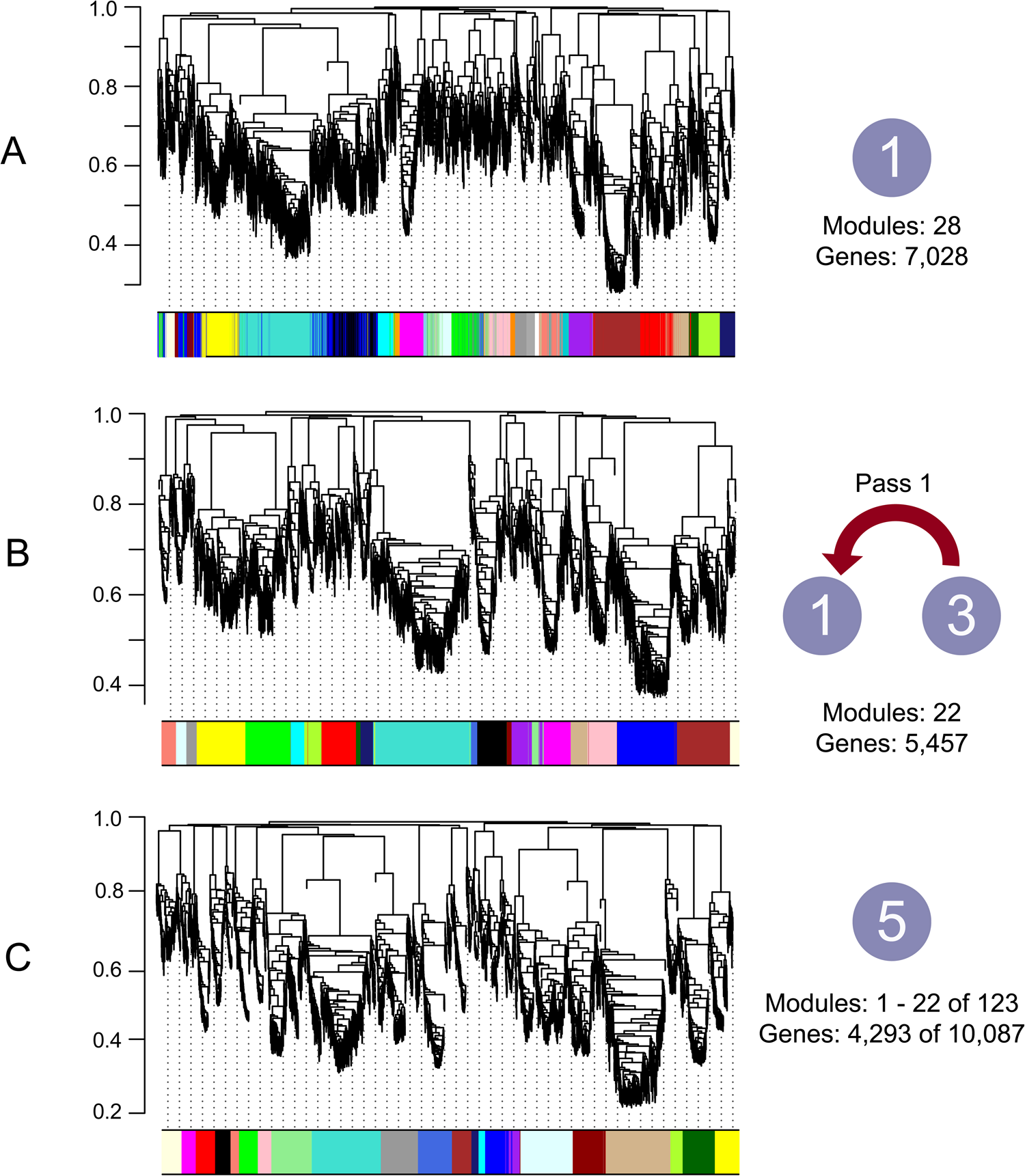
Iterative refinement of co-expression networks improves WGCNA clustering of whole transcriptome gene profiling. Hierarchical clustering of genes generated from blockwise and iteratively refined WGCNA analyses of the 12,397 genes expressed in the ADULT dataset. The colored strips under each dendrogram depict the module assignment for the gene associated with the corresponding dendrogram branch. Module colors are not consistent among panels. Large, blue numbered circles refer to steps in the refined WGCNA analysis outlined in **Fig. 2**. A. Clustering and detected modules resulting from blockwise WGCNA analysis (result after Step 1-2; **Fig. 2**), with poorly fitting genes removed yields a cleaner result, but module genes are still distributed across multiple clusters. This filtering also has the added consequence of significant information loss (>45% of genes pruned). B. Result from the first pass of iterative refinement of WGCNA: repeat steps 1-3 (**Fig. 2**) until no more genes are pruned. This refinement markedly improves the clustering of genes fitting the primary signal in the expression dataset. Gene clusters are distinct (separated by long branches) and only a handful of module members are distributed across multiple clusters. However, there is significant additional information loss compared to A. C. Same core modules illustrated in A and B, after final network assembly (step 5; **Fig. 2**). Clustering is further refined compared to the first pass (B, above); no modules are distributed across multiple clusters. In addition to the core set of 22 modules illustrated here (blocks along the diagonal in **Fig. 1E**), iterativeWGCNA was able to detect an additional 100 modules, each of which groups coherently expressed genes, within the overlapping network sub-structure (not figured). Thus, by reprocessing the pruned genes, this approach was able to improve not only the quality of the detected modules, but also classify >3000 more genes compared to standard blockwise WGCNA analysis (A, above).

IterativeWGCNA reapplies the *blockwiseModules* function to the gene expression dataset retained after pruning. In doing so, each iteration of iterativeWGCNA there is a recalculation of topological overlap, recalculation of the eigengenes, and a redefinition and reassessment of gene module membership. The process is unsupervised, repeating itself as many times as necessary for the network structure to stabilize, ultimately producing a cleaner, more refined clustering and classification of the GCN (**Fig. 3B**).

Although this refining improves the resolution of the GCN, there is a trade-off: better clusters are obtained at the expense of information loss. In the case of the ADULT dataset, the initial filtering done by WGCNA’s *blockwiseModules* resulted in loss of >45% of the expressed genes in the dataset (**Fig. 3A**). A single pass of iterativeWGCNA yield resulted in a refined GCN, but a loss of ~1,500 additional genes, increasing the total information loss to ~58% of the gene set. These genes do not fall into the “garbage bin” category pruned during the first application of the *blockwiseModules* function; they do not have variable or ubiquitous expression but were instead simply misplaced by the greedy hierarchical clustering. To bring as many of these genes back in, and find a globally optimal module assignment for them, iterativeWGCNA makes a new pass at the data, this time with the residuals to the fit (all genes pruned in that initial pass; step 4 **Fig. 2**) and again iterating until no new modules are detected. The entire process is repeated until no new modules are detected from the residual gene sets. The end result is a cleaner and more inclusive result; in the case of the ADULT dataset, applying iterativeWGCNA allows >75% of the genes to be classified (**Fig. 3C**).

The effectiveness of iterativeWGCNA is illustrated in **Fig. 4**, which tracks 396 genes assigned to a “core” module detected by WGCNA in the ADULT dataset through the various iterations and passes of iterativeWGCNA. Module 6 (WGCNA M6) was detected after the first application of *blockwiseWGCNA* to the ADULT dataset; red **Fig. 4A**). This is a fairly robust module; 70% of the genes assigned to this cluster form its central core and stay together for the duration of the iterative analysis. By the end of the final iteration (I9) of first pass (P1), iterativeWGCNA has refined the analysis, pruning the 99 additional genes and reassigning 114 different genes initially assigned to another module (M12) or left unclassified, as consequence of changes in the network topology (**Fig. 4A**). Although the two modules are of similar orders of magnitude and essentially defined by the same eigengene (**Fig. 4C**), the one detected by a refining pass of iterativeWGCNA, P1_I9_M3 (n = 390 genes; dark green, **Fig. 4**), comprises genes with more coherently expressed than what was originally detected by WGCNA (M6; **Fig. 4B**).

**Figure 4.**
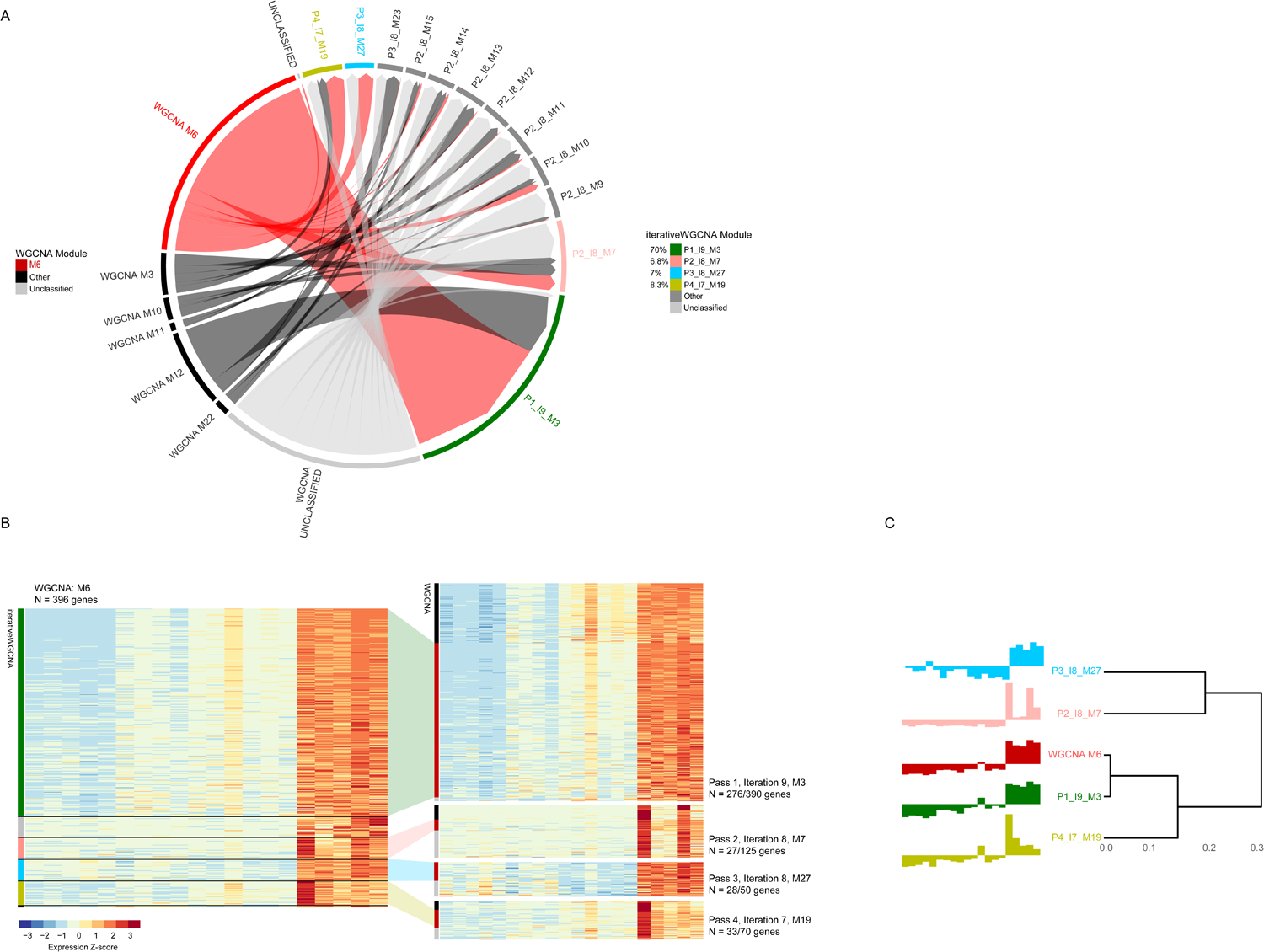
Information gain and module refinement via application of the iterativeWGCNA method is illustrated by tracking changes in module membership of genes assigned to a single module by WGCNA. A. Chord diagram illustrating how gene membership changes through each step of the iterativeWGCNA process. Modules on the left side of the chord diagram were detected by blockwise WGCNA; to the right is the final module membership after applying iterativeWGCNA to the same expression dataset. 396 genes were assigned by blockwise WGCNA (after pruning) to module 6 (WGCNA M6; red). To account for changes in network topology, WGCNA was reapplied (steps 1-3; **Fig. 2**) until the network stabilized; 9 iterations in total. By this point – pass 1 (P1), iteration 9 (I9) – the module membership had been refined (P1_I9_M3, dark green) due to both the removal of outliers removed and the incorporation of better-fitting genes from other modules (black) or the unclassified set (grey) pulled in. The process was repeated on the residual gene set (step 4 and repeat 1-3; **Fig. 2**) and allowing detection of new modules from the unclassified gene set and better placement of genes that were poor fits to their WGCNA-assigned module. B. Standardized expression heatmaps illustrating improved grouping of coherently expressed genes using iterativeWGCNA compared to standard blockwise WGCNA analysis. To the left is the expression heatmap for WGCNA M6 (red in A, above). Colored indicators on the left border of the heatmap indicate final module assignment after applying iterativeWGCNA (colors as in A, above). 92.2% of the genes were reassigned to just 4 modules: 70% to P1_I9_M3 (dark green), 6.8% to P2_I8_M7 (pink), 7% to P3_I8_M27, and 8.3% to P4_I7_M19. Expression heatmaps for these modules are drawn on the right; colored indicators on the left border indicate original WGCNA module assignment (M6: red, other: black; unclassified: grey). C. Dendrogram illustrating the eigengene similarity of the WGCNA M6 module to the four modules detected by iterativeWGCNA and highlighted in B, left.

Over the course of subsequent passes of iterativeWGCNA, the 30% remaining WGCNA M6 member genes are split among modules detected among the residual genes. Like P1_I9_M3, these modules are comprised of genes that were either initially left unclassified by WGCNA (often >50% of module membership; **Fig. 4A**), or assigned to their first fit, but not best, module. Of these, 3 modules stand out, together containing 22.1% of the non-core genes from WGCNA M6 (red; **Fig. 4**). These are 1) P2_I8_M7 (pink; detected in the 8^th^ iteration of the second pass), containing 6.8% of the WGCNA M6 genes, 2) P3_I8_M27 (blue); 7%, and 3) P4_I7_M19 (yellow ochre); 8.3%. The eigengenes of these modules are high correlated (*r*^2^>0.8; **Fig. 4C**), but most definitely distinct; grouping distinct gene sets (**Fig. 4B**).

## Discussion

Here we have introduced an iterative-extension to the popular WGCNA package that minimizes information loss and provides an unbiased, unsupervised means for identifying coherently expressed gene sets in transcriptome-wide expression datasets. The method works by recognizing that most GCNs possess a central “core” defining the broader structure of the GCN. A standard WGCNA result is delimited by the strong signal established by these core gene sets and the greedy nature of hierarchical clustering, which prioritizes assigning genes to existing clusters over creating new ones. The more a network deviates from this central core (e.g., **Fig. 1E**), the harder it is to impose a scale-free topology and the more poorly WGCNA will perform.

Pruning poorly fitting genes improves the overall look of a WGCNA-based classification; however, it also can dramatically alter the network architecture, in terms of both modularity and topological overlap by reducing the number of inter-modular connections in the GCN. Our approach accounts for this by iteratively recalculating and re-clustering the GCN, reassessing gene module memberships until the network structure stabilizes. The GCNs inferred from all three datasets considered in this study do approach scale-free topology after power-law weighting and as such, exhibit fairly stable network modularity throughout each iteration, despite changes in topological overlap. As a supplement to this analysis and to better illustrate the impact of pruning on network topology, we are currently investigating a published RNA-seq expression dataset of similar magnitude for which the scale-free topology fit index does not approach values >0.8 for reasonable power levels (< 30).

Although iterativeWGCNA marks an improvement over standard WGCNA analysis by reducing overall information loss while improving the modularity of detected gene sets, there is a trade-off in the total number of detected modules. IterativeWGCNA frequently detects several times more modules that standard WGCNA analysis (e.g. 28 compared to >100 for the ADULT dataset). It raises the question as to whether the approach over splits the data and how much that splitting depends on parameter choices, namely the eigengene similarity cut-off. We plan to illustrate that these clusters are biologically meaningful via consensus network analysis of the FETAL and ADULT datasets, to show that the method successfully prunes noisy data, while isolating contaminant gene signatures and key differences in gene-expression between adult and fetal definitive erythropoiesis.

## Conclusions

We have described the iterative application of a WGCNA-based strategy to derive a robust gene co-expression networks for whole-transcriptome datasets. This improved computational strategy greatly expands the general applicability of WCGNA and provides a robust and hierarchical framework for deriving and exploring GCNs at increasing levels of resolution and heterogenous conditions.

## Acknowledgements

This study was supported by funding from the NIH to MAM (DK72473 and DK89523).

## References

Gautier, L., 2012. rpy2: A simple and efficient access to R from Python. https://doi.org/ https://rpy2.bitbucket.io/

Gautier, L., Cope, L., Bolstad, B.M., Irizarry, R.A., 2004. affy—analysis of Affymetrix GeneChip data at the probe level. Bioinformatics 20, 307–315. https://doi.org/10.1093/bioinformatics/btg405

Horvath, S., 2017. WGCNA package: Frequently Asked Questions [WWW Document]. URL https://labs.genetics.ucla.edu/horvath/CoexpressionNetwork/Rpackages/WGCNA/faq.html (accessed 11.20.17).

Kingsley, P.D., Greenfest-Allen, E., Frame, J.M., Bushnell, T.P., Malik, J., McGrath, K.E., Stoeckert, C.J., Palis, J., 2013. Ontogeny of erythroid gene expression. Blood 121, e5–e13. https://doi.org/10.1182/blood-2012-04-422394

Langfelder, P., Horvath, S., 2008. WGCNA: an R package for weighted correlation network analysis. BMC Bioinformatics 9, 559. https://doi.org/10.1186/1471-2105-9-559

Massa, A.N., Childs, K.L., Lin, H., Bryan, G.J., Giuliano, G., Buell, C.R., 2011. The Transcriptome of the Reference Potato Genome Solanum tuberosum Group Phureja Clone DM1-3 516R44. PLOS ONE 6, e26801. https://doi.org/10.1371/journal.pone.0026801

R Core Development Team, 2017. R: A Language and Environment for Statistical Computing. R Foundation for Statistical Computing, Vienna, Austria.

Wu, J., Gentry, R.I. with contributions from J.M.J., 2017. gcrma: Background Adjustment Using Sequence Information.

Xue, Z., Huang, K., Cai, C., Cai, L., Jiang, C., Feng, Y., Liu, Z., Zeng, Q., Cheng, L., Sun, Y.E., Liu, J., Horvath, S., Fan, G., 2013. Genetic programs in human and mouse early embryos revealed by single-cell RNA sequencing. Nature 500, 593–597. https://doi.org/10.1038/nature12364

Zhang, B., Horvath, S., 2005. A General Framework for Weighted Gene Co-Expression Network Analysis. Stat. Appl. Genet. Mol. Biol. 4. https://doi.org/10.2202/1544-6115.1128

